# Enterovirus D68 VP1 and VP3 determine neurotropism in human spinal cord organoids

**DOI:** 10.64898/2026.01.25.701582

**Authors:** Jessica E Packard, Jennifer E Jones, Gal Yovel, Megan Culler Freeman

**Affiliations:** Department of Pediatrics, University of Pittsburgh School of Medicine, Pittsburgh, PA, USA; Institute of Infection, Inflammation, and Immunity in Children (i4Kids), Pittsburgh, PA, USA

## Abstract

Enterovirus D68 (EV-D68) is a non-polio enterovirus that can cause a polio-like paralysis condition, acute flaccid myelitis (AFM). EV-D68 associated AFM cases waned in the US after 2018 and the reasons for this are unknown. It has recently been demonstrated that EV-D68 containing point mutations in viral structural proteins VP1 and VP3 resulted in decreased paralysis in different neonatal mouse models. However, phenotypes of these mutations in a human multicellular central nervous system (CNS) model are unknown. We hypothesized that mutations in VP1 and VP3 will similarly direct neurotropism in human spinal cord organoids (hSCO). To investigate this, we recreated viruses with mutations in VP3 (I88V) or VP1 (L1I/N2D/T98A/E283K or L1P/V148A/K282R) and infected hSCOs. We found that VP3 I88V and VP1 L1I/N2D/T98A/E283K resulted in decreased titer and viral protein staining, consistent with attenuated neurovirulence in previously published murine models. When these mutations were combined, their effects on neurotropism were not additive. Sequence analysis of recently circulating EV-D68 strains reveals that VP3 I88 and VP1 E283 have remained the dominant amino acid residues since 2014, whereas VP1 sites 1, 2, and 98 have higher population diversity, indicating that these residues may be contributing to newly reduced neurovirulence after 2018.

## INTRODUCTION

Enterovirus D68 (EV-D68) is a non-enveloped enterovirus within the *Picornaviridae* family originally isolated from patient respiratory samples in 1962 [1]. Today, EV-D68 is considered a reemerging virus due to increased circulation, severe respiratory illness and new neurovirulence. EV-D68 infections are linked to acute flaccid myelitis (AFM), a polio-like illness resulting in paralysis, primarily in children [1, 2].

Prior studies investigating EV-D68 determinants of neuropathogenesis have used mouse models to study the relationship between EV-D68 infection and the development of paralysis. Two groups identified mutations in structural proteins VP1 and VP3 as primary drivers of neuropathogenesis [3, 4]. Yeh et al. 2020 found that in EV-D68 US/IL/14-18952 (EV-D68 18952), introduction of a mutation in VP3 at site 88 from isoleucine to valine (VP3 I88V), as present in MO/14-18949, resulted in decreased paralysis and viral titer in day seven C57BL/6-derived Tg21 mice with type I interferon receptor knockout (Tg21/IFNR-ko) after intraperitoneal (IP) injection [3]. The authors found that five day old C57BL/6-derived Tg21 mice infected with VP3 I88V via intracranial (IC) injection had significantly decreased viral titer in the brain and spinal cord compared to EV-D68 18952. Additional VP1 mutations, L1P, V148A, K282R, and G283E were investigated. They found that VP1 mutations in L1P and K282R/G283E by IP injection resulted in decreased viral titer and paralysis in seven-day old mice at 24-hours post infection (hpi) and 72 hpi.

Leser et al. 2024 also sought to evaluate EV-D68 neuropathogenesis determinants by comparing neurovirulent EV-D68 18952 and avirulent CA/14-4231 (CA4231) [4]. They found that while EV-D68 18952 results in paralysis after intramuscular (IM) injection in Swiss Webster interferon deficient (mitochondrial antiviral signaling protein (MAVS^−/−^)) mice bred on a pure C57BL/6 background, CA4231 does not. Through systematic evaluation of the differences between the strains, they discovered that four VP1 amino acid mutations present in CA4231 (L1I/N2D/T98A/E283K) were the primary drivers of EV-D68 neurovirulence.

The discordance between the mutations responsible for driving neuropathogenesis may be reconcilable for a few reasons [3, 4]. Primary differences between the two studies include the use of mouse genetic backgrounds, inoculation route used (IM vs IP vs IC injection), and mapping to different avirulent strains (MO vs. CA). To assess EV-D68 determinants of neurovirulence independent of these variables, we sought to investigate if these same mutations conferred a similar phenotype in a human multicellular model with intact elements of innate immunity [5]. Specifically, we generated recombinant EV-D68 (rEV-D68) strains to match the VP3 mutation from Yeh et al. (rVP3-I88V-MO), three amino acid changes in VP1 described by Yeh et al. (rVP1 3AA-MO), and four amino acid changes described by Lesser et al. (rVP1-4AA-CA).

Using a previously established multicellular human spinal cord organoid (hSCO) model for EV-D68 neurotropism, we evaluated the replication competence of each rEV-D68. All rEV-D68 containing mutations replicated in our hSCO model to differing degrees. Both rVP3 I88V-MO (strain from Yeh et al. [3]) and VP1 4AA-CA (strain from Leser et al. [4]) produced significantly decreased viral titer in hSCOs at 48 and 96 hpi as compared to the reference strain, consistent with phenotypic conclusions from the murine models. These findings suggest that while these viruses may have potential defects in accessing the CNS from the initial infection site in murine models, they also have decreased replication capacity in CNS cells. Because the viral strains in Yeh and Leser were from 2014 and prior, we sought to evaluate the residues of interest in modern circulating populations by assessing mutational frequency over time. We found that VP3 I88 and VP1 E283 have been relatively fixed in viral populations since 2014, however, there has been more population level diversity at VP1 positions 1, 2, and 98. As clinical AFM attributed to EV-D68 has waned since 2018, these residues may be driving reduced neurovirulence.

## MATERIALS AND METHODS

### Cells and Viruses

Human embryonic kidney (HEK) 293T cells (ATCC CRL-3216) and human RD cells (ATCC CCL-136) were obtained from ATCC. Human induced pluripotent stem cells (iPSC, SCTi003-A) were obtained from STEMCELL Technologies. HEK 293T and RD cells were maintained in Dulbecco’s Modified Eagle’s Medium (DMEM, Corning) supplemented with 10% fetal bovine serum (FBS, Biowest) and 1% penicillin/streptomycin (Gibco). Human iPSCs were maintained in mTeSR^TM^ Plus medium (STEMCELL Technologies) or Essential 8 medium (Thermofisher) in flasks coated with 150 µg/mL Cultrex (R&D Systems). Cells were maintained in sterile cell culture incubators at 37°C, 5% CO_2_ unless otherwise stated.

### Cloning and generation of rEV-D68

EV-D68 infectious clones were obtained through BEI Resources, National Institute of Allergy and Infectious Diseases (NIAID), National Institute of Health (rUSA/IL/2014-18952 from pUC19-EVD68_49131 (NR-52011) and rUSA/Fermon from pUC19-R-Fermon) (NR-52375). Point mutations were inserted into the pUC19-EVD68_49131 using the NEBuilder HiFi DNA Assembly Method based on previously published literature [6]. To generate the VP3 mutation, the codon for isoleucine at site 88 was replaced with a valine (I88V) (rVP3-I88V-MO) (Table 1). To replicate the mutations from Yeh et al. in VP1, we altered three amino acids (rVP1-3AA-MO): leucine at site 1 to proline (L1P), valine at site 148 to alanine (V148A), and lysine at site 282 to arginine (K282R). Yeh et al. also identified a fourth point mutation at site 283 in VP1 (glycine to glutamic acid at site 283 (G283E)) [3]. However, our EV-D68 18952 strain already encodes E283, therefore we did not make a point mutation at VP1 site 283. To generate four amino acid mutations from Leser et al. in VP1 (rVP1-4AA-CA), the following changes were made: leucine at site 1 to an isoleucine (L1I), asparagine at site 2 to aspartic acid (N2D), threonine at site 98 to an alanine (T98A), and glutamic acid at site 283 to a lysine (E283K). To generate rVP3-I88V/VP1-4AA-CA combined mutations, we made five-point mutations, one in VP3 (I88V) and four in VP1 (L1I/N2D/T98A/E283K). PCR primers were created according to Table 1.

**Table 1.**
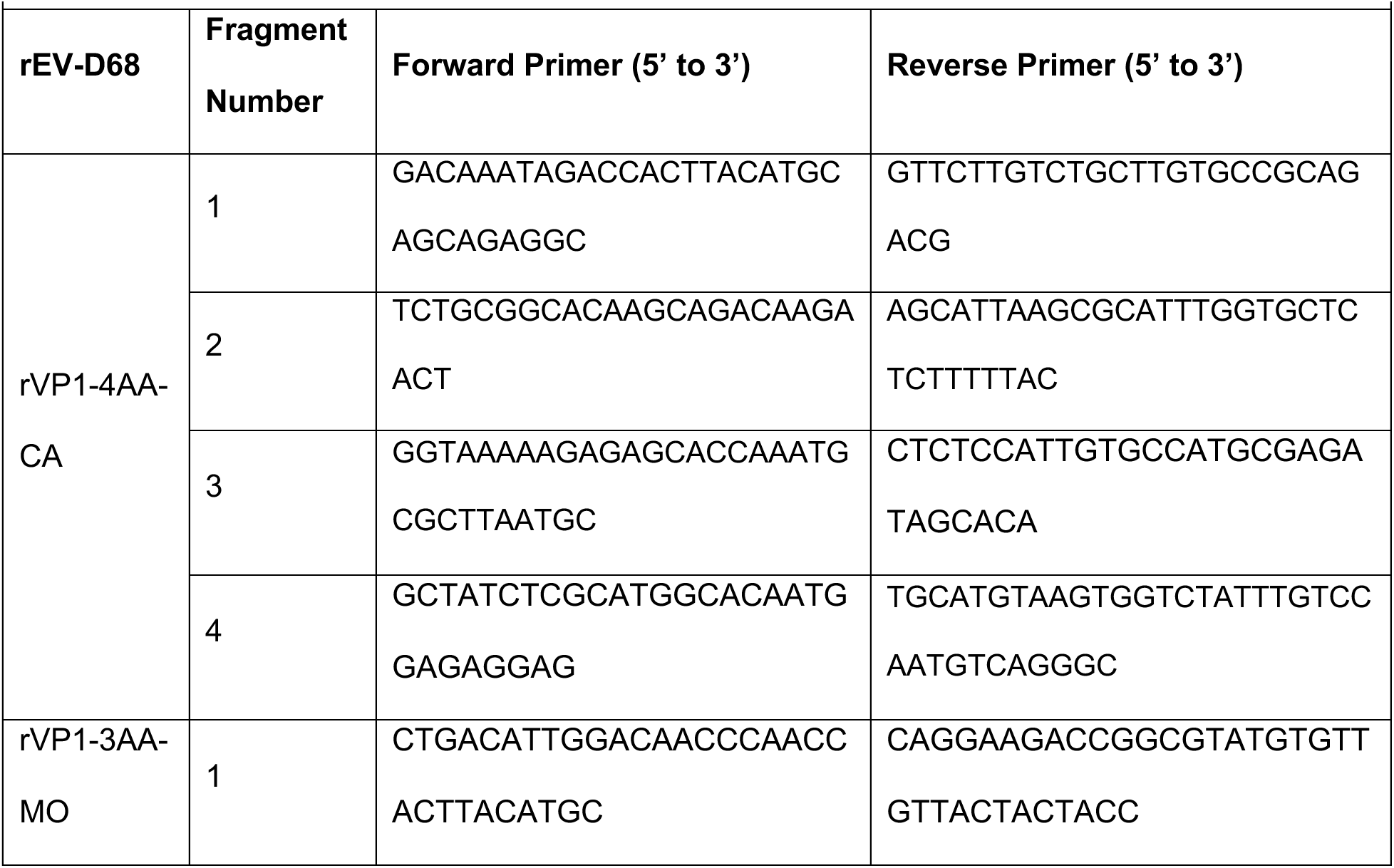
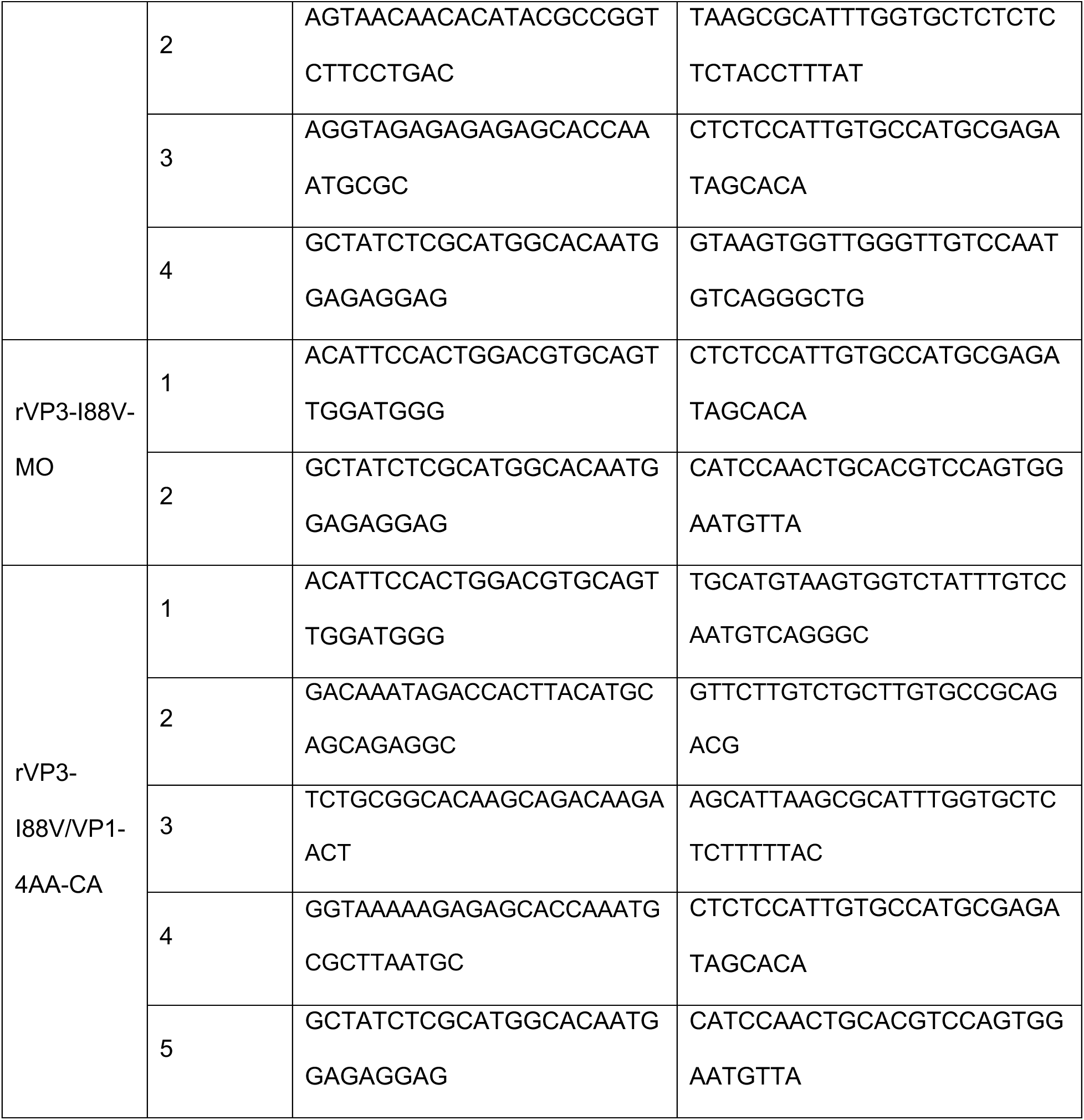
Primers used for NEBBuilder HiFi DNA assembly to generate VP3 and VP1 recombinant EV-D68 (rEV-D68).

DNA was amplified by PCR and run on a 1% agarose gel. Gel fragments were cut according to size and DNA extracted using the Thermofisher Scientific GeneJET Gel Extraction Kit. For the VP3 point mutation, two fragment assembly was used with a vector to insert ratio of 1:2. For VP1 three and four-residue mutations, four fragment assembly was used with a vector to insert ratio of 1:1. For the rVP3-I88V/VP1-4AA-CA five-residue mutation, five fragment assembly was used with a vector insert ratio of 1:1.The combined PCR products were digested with DpnI to destroy the plasmid template before setting up the assembly reaction. Samples were assembled by a 60-minute incubation at 50°C. Assembled product (2µL) was transformed into NEB 10-beta competent *E. coli* cells.

Following successful transformation, plasmids were extracted using GeneJet Mini Plasmid Prep Kit according to the manufacturer’s protocol. To verify successful insertion of viral mutations, whole genome sequencing was performed using long-read sequencing technology from Oxford Nanopore Technologies by Plasmidsaurus and was compared to EV-D68 18952.

### Transfection and Virus Rescue

HEK293T cells were seeded into 100 mm^2^ dishes. Plasmids were transfected with pCAGGS T7 at a ratio of 10:1 with the Transit-LT1 transfection reagent (Mirus Bio) into HEK293T cells. Cells were incubated at 33°C, 5% CO2 until cytopathic effects were visible. Cells were scraped from the dish and collected with supernatant.

Viral stocks were obtained by incubation of transfected lysates on RD cells. RD cells were seeded with 20 million cells in 150 mm^2^ dishes. Half of the transfection lysate was added to RD cells and rocked at room temperature for one hour to adsorb virus. Mock infected cells received cell culture media only. Inoculum was aspirated from cells and fresh cell growth media was added. Cells were incubated at 33°C, 5% CO_2_ until cytopathic effects. Viral stocks were harvested and purified using sucrose cushion centrifugation as previously described [7].

### Growth kinetics in human cell lines

RD cells were seeded at 200,000 cells per well and grown in a 24-well tissue culture plate. RD cells were infected with rescued recombinant virus at an MOI of 0.01 plaque forming units (PFU)/cell. Cells were inoculated in biological triplicate with the indicated recombinant virus or mock-infected with PBS. Virus was adsorbed onto cells for one hour at room temperature with rocking. Following one hour, the inoculum was aspirated, cells were washed three times with PBS, and fresh growth medium was added to cells. RD cells were incubated for 96 hours at 33°C, 5% CO_2_. Supernatants were taken at 0, 24, 48, 72, and 96 hpi and stored at -80°C. Infectious virus was quantified by 50% tissue culture infectious dose assay (TCID_50_) on RD cells in technical triplicate. TCID_50_ titers were enumerated by the Spearman-Kärber method.

### Differentiation and propagation of human spinal cord organoids

3-dimensional spinal cord (3-DiSC) organoids were differentiated from iPSC cells as previously described [5]. iPSC cells were seeded at a density of 9,000 cells per well into a 96-well round-bottom low adhesion plates using Accumax (Sigma). Cells were plated in differentiation medium for 14 days supplemented with designated growth factors [5]. After 14 days, organoids were transferred to a tissue culture plate and cultured in suspension in N2B27 medium [DMEM/F-12 (Gibco), neurobasal medium (Gibco) (1:1), 0.5% (vol/vol) N2 supplement (ThermoFisher) and 1% (vol/vol) B27 supplement without vitamin A (Gibco)] supplemented with 1 mM L-glutamine (ThermoFisher), 0.1 mM β-mercaptoethanol, 0.5 µM ascorbic acid, 10 ng/mL brain-derived neurotrophic factor (BDNF, STEMCELL Technologies), 10 ng/mL glial cell line-derived neurotrophic factor (GDNF, STEMCELLL Technologies), and 100 nM retinoic acid (Tocris)).

### Growth kinetics in human spinal cord organoids

Pools of 12 3-DiSC hSCOs were inoculated with rescued recombinant virus in biological triplicate at 14 days post-propagation. hSCOs were infected with 10^4^ PFU per pool in N2B27 medium. Pools were either infected with the indicated recombinant virus or with a mock inoculum of PBS. Virus was adsorbed on hSCOs for one hour at room temperature with rocking. After one hour, inoculum was aspirated, and organoids were washed three times with PBS. Pools of hSCOs were then transferred to new plastic to ensure there was no residual inoculum present and fresh growth medium was added. hSCOs were incubated for 96 hours at 33°C, 5% CO_2_. Supernatants were taken at 0, 24, 48, 72, and 96 hpi and stored at -80°C. At 96 hpi, hSCO were fixed with 4% paraformaldehyde in PBS for IF staining. Infectious virus was quantified by 50% tissue culture infectious does assay (TCID_50_) on RD cells in technical triplicate. TCID_50_ titers were enumerated by the Spearman-Kärber method.

### Immunofluorescence staining

IF staining on fixed hSCOs was completed at previously described [5]. hSCOs were washed with 1% PBS-BSA and PBS-Tween at 4°C. The remaining washes and antibody dilutions were diluted in organoid wash buffer (0.1% Triton X-100 (v/v), 0.2% BSA (m/v) in PBS). Primary antibodies VP1 (Genetex, GTX132313) and Phalloidin stain 543 (Biotium, 00043-T) were used at a 1:250 dilution. Secondary antibody, Alexa Fluor^TM^ 488 (A-11001), was used at a 1:1000 dilution. DAPI (4’,6-diamidino-2-phenylindole, dihydrochloride) (ThermoFisher, D1306) was used at a 1:300 dilution. hSCOs were cleared for 30 minutes at with 60% glycerol in 2.5 M fructose before mounting and imaging.

### Confocal imaging

Microscope slides were imaged on a Leica Stellaris 5 confocal microscope. Volumetric Z-stacks were imaged with a 20X objective by oil immersion. Sequential scanning was used with 2X line averaging by frame. The following parameters were used: dsDNA was measured at 403 nm (4’,6-diamidino-2-phenylindole (DAPI)), Phalloidin stain was measured at 543 nm, and VP1 was measured at 488 nm. Images were processed with FIJI 2 version 2.9.0.

### Statistical analysis

All statistical analyses were performed using Graphpad Prism version 10. The statistical test used for each figure is indicated in the figure legend. All experiments were performed in at least biological triplicate corresponding to each experimental condition. Each biological replicate consisted of an average of three technical replicates.

### Analysis of substitution frequencies

EV-D68 FASTA sequences (n=7,468) were downloaded from the Bacterial and Viral Bioinformatics Resource Center (BV-BRC, accessed November 26, 2025). Partial sequences or sequences isolated outside of the 2014-2024 timespan were excluded, leaving 1,968 for analysis. Sequences were aligned in NextClade (https://clades.nextstrain.org) [8] using the EV-D68 dataset with reference Fermon (accession no. AY426531.1) [9] as a basis for QC, clade assignment, and mutation calling. Sequences that failed QC were excluded (n=2). Data were visualized in R (version 4.4.1) using the expss (version 0.11.7) and areaplot (version 2.1.3) packages.

## RESULTS

### Recombinant EV-D68 with mutations in VP1 and VP3 replicate efficiently in RD cells

We recreated three rEV-D68 viruses as previously described in the EV-D68 18952 backbone : one rEV-D68 with four amino acid changes in VP1 (designated rVP1-4AA-CA), one rEV-D68 with three amino acid changes in VP1 (designated rVP1-3AA-MO) and one rEV-D68 with a point mutation in VP3 (designated rVP3-I88V-MO) (Figure 1A) [3, 4]. We then inserted point mutation fragments into the pUC19-EVD68_49131 vector using the NEBuilder HiFi DNA Assembly Method [6]. As controls, we also created plasmids expressing EV-D68 US/IL/14-18952 (designated rEV-D68 18952) and EV-D68 Fermon (designated rFermon) [5]. We then transfected plasmids into HEK293T cells with pCAGGS T7 at a ratio of 10:1 with the Transit-LT1 transfection reagent (Figure 1B). After observable cytopathic effects (CPE), we collected supernatants and created viral stocks.

**Figure 1.**
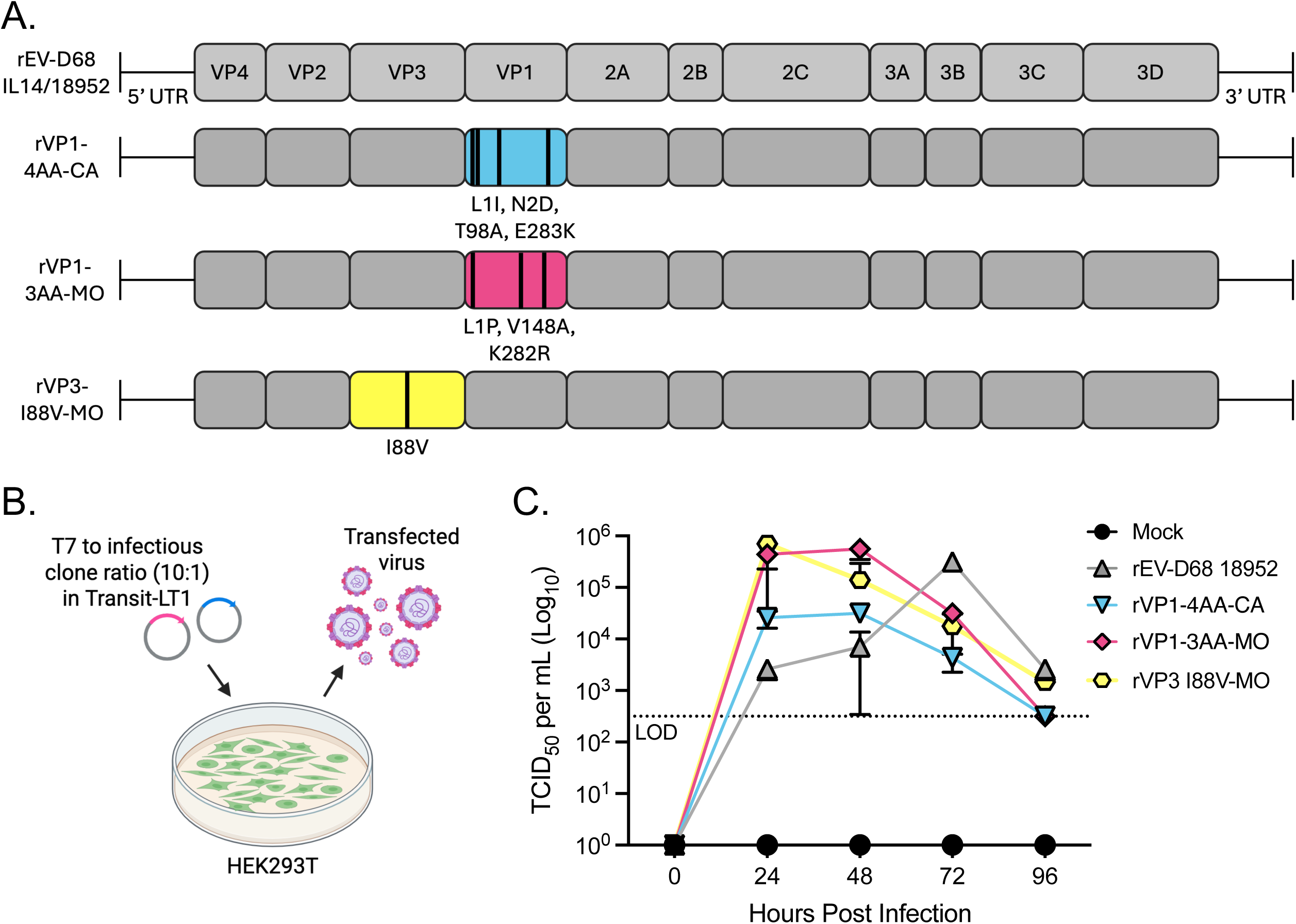
Confirmation of recombinant EV-D68 with VP1 and VP3 mutations. A.) Schema of proposed recombinant viruses with point mutations in either VP1 or VP3. Gray represents rEV-D68 18952, blue – rVP1-4AA-CA (L1I/N2D/T98A/E283K), pink – rVP1-3AA-MO (L1P/V148A/K282R), and yellow – rVP3-I88V-MO. B.) Infectious clones were cloned into pUC19-EV-D68_49131 and were transfected with pCAGGS Y7 at a ratio of 10:1 with the Transit-LT1 transfection reagent (Mirus Bio) into HEK293T cells. Cells were incubated until cytopathic effects were visible and then were collected. Viral stocks were obtained by incubation of transfected lysates on RD cells. Created with Biorender.com. Schematic is modeled from Jones et al. 2024 [6]. C.) RD cells were infected at an MOI 0.01 PFU/cell in biological triplicate with each recombinant virus. Viral titers were determined by TCID_50_ in RD cells. Black circle – mock, gray triangle – rEV-D68 18952, blue upside-down triangle – rVP1-4AA-CA, pink diamond – rVP1-3AA-MO, yellow hexagon – rVP3-I88V-MO.

To determine the replication competence of recovered rEV-D68s, RD cells were infected with each rEV-D68 at an MOI 0.01 PFU/cell. Supernatants were collected between 0- and 96-hpi at 24-hour intervals followed by titer determination by TCID_50_. We found that infectious virus was detectable at 24 hpi for all recombinant viruses (Figure 1C). These data indicate that we successfully rescued replication-competent recombinant EV-D68 without growth defects in RD cells.

### rVP3-I88V-MO and rVP1-4AA-CA have decreased viral replication in hSCOs

We next wanted to determine if point mutations VP1 or VP3 influence neurotropism in our hSCO model. hSCOs were mock-infected or infected with 10^4^ PFU rEV-D68. We collected infected hSCOs at 96 hpi and stained hSCOs for VP1, an EV-D68 structural protein, and phalloidin, a filamentous actin stain (Figure 2A). We then quantified the mean fluorescence intensity (MFI) of VP1 stain across all hSCOs (Figure 2B). We found that there was little VP1 staining in mock- or rFermon-infected hSCOs, consistent with previously published results using the clinical strain [5]. We also found that compared to rEV-D68 18952, there was a significant decrease in VP1 staining in rVP3-I88V-MO and rVP1-4AA-CA.

**Figure 2.**
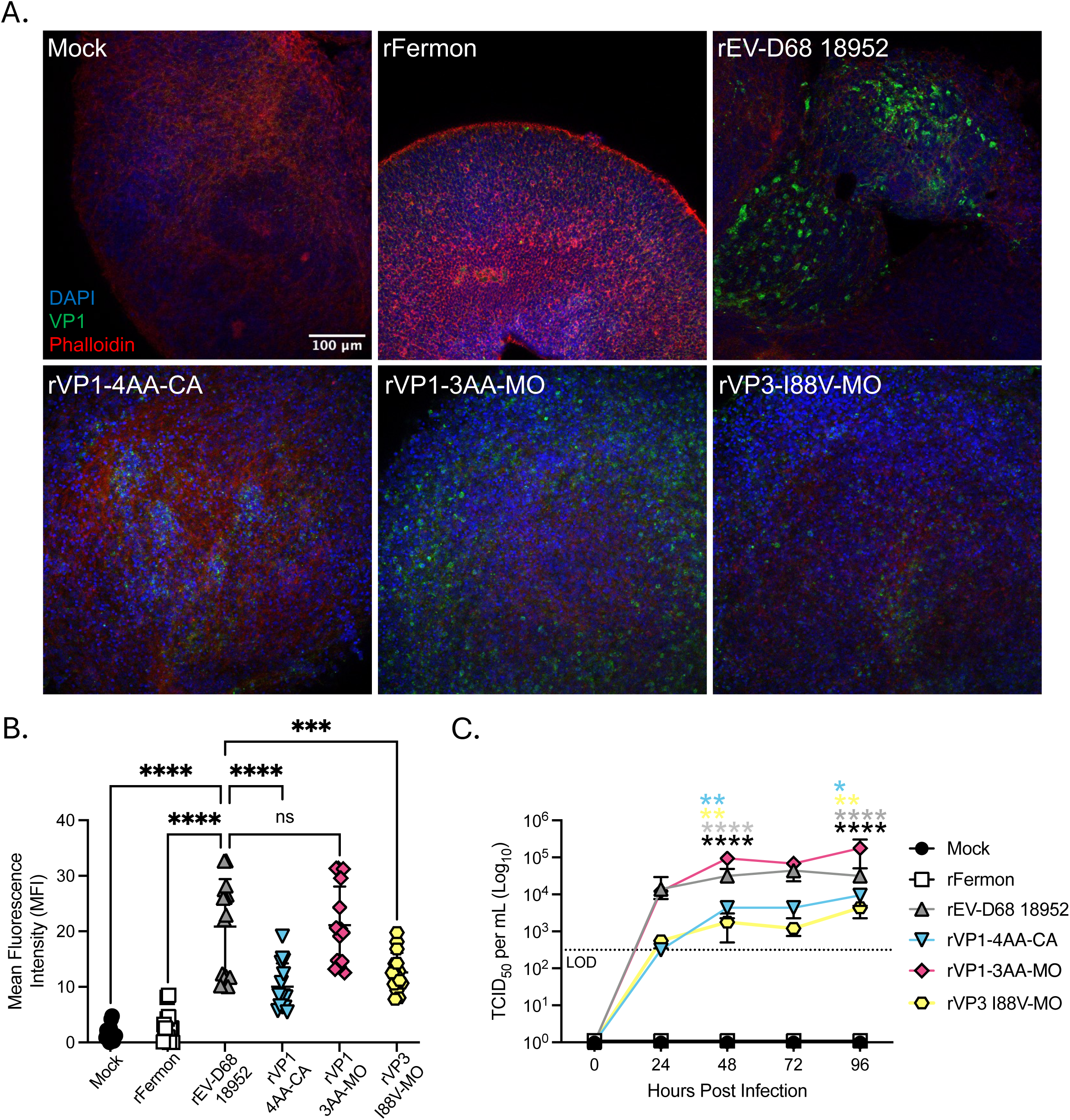
rVP1-4AA-CA and rVP3-I88V-MO result in decreased infection of hSCOs. A.) IF images of SCTi003A hSCOs infected with recombinant viruses. hSCOs were infected with 10^4^ PFU and were fixed in 4% paraformaldehyde at 96 hpi. hSCOs were then stained with DAPI (blue), VP1 (green), and phalloidin (red). B.) MFI of VP1 staining was quantified across five biological replicate images of each recombinant virus across multiple Z stacks and organoids. Quantified values were calculated using Fiji ImageJ. One way ANOVA with multiple comparisons to EV-D68 18952 was performed (ns – no significance, *** p < 0.001, **** p < 0.0001). C.) SCTi003A hSCO were infected with 10^4^ PFU of each recombinant virus in biological triplicate. Supernatants were collected every 24 hours up to 96 hpi. Viral titers were determined by TCID_50_ in RD cells. A two-way ANOVA with multiple comparisons to EV-D68 18952 was performed (* p < 0.05 ** < 0.005 **** < 0.00005). Black asterisks – compared to Mock, gray asterisks – compared to rFermon, blue asterisks – compared to rVP1-4AA-CA, and yellow asterisks – compared to rVP3-I88V-MO. Black circle – mock, white square – rFermon, gray triangle - rEV-D68 18952, blue upside-down triangle – rVP1-4AA-CA (L1I/N2D/T98A/E283K), pink diamond – rVP1-3AA-MO (L1P/V148A/K282R), and yellow hexagon – rVP3-I88V-MO.

As a complementary approach, we then assessed rEV-D68 growth in hSCO. hSCOs were pooled as described previously and organoids were infected with 10^4^ PFU per pool (Figure 2C). We found that rFermon did not produce measurable titer in hSCOs consistent with previously published data using the clinical strain [5]. At 48 and 96 hpi, rVP3-I88V-MO and rVP1-4AA-CA resulted in decreased viral titer compared to control strain, rEV-D68 18952. These data suggest that rVP3-I88V-MO and rVP1-4AA-CA, but not rVP1-3AA-MO, have decreased fitness in cells of the human CNS.

### rVP3-I88V/VP1-4AA-CA results in decreased viral replication in hSCOs compared to rEV-D68 18952

After determining that both rVP3-I88V-MO and rVP1-4AA-CA resulted in decreased viral titer and VP1 expression, we wanted to determine if combining all five mutations into a single recombinant virus would have an additive effect on neurotropism. We inserted VP3 I88V and VP1 L1I, N2D, T98A, and E283K point mutation fragments (designated rVP3-I88V/VP1-4AA-CA) into the pUC19-EVD68_49131 vector (Figure 3A) as previously described followed by transfection into HEK293T cells [6]. We successfully recovered this rEV-D68 and rVP3-I88V/VP1-4AA-CA replicated efficiently in RD cells between 0 and 96-hpi (Figure 3B).

**Figure 3.**
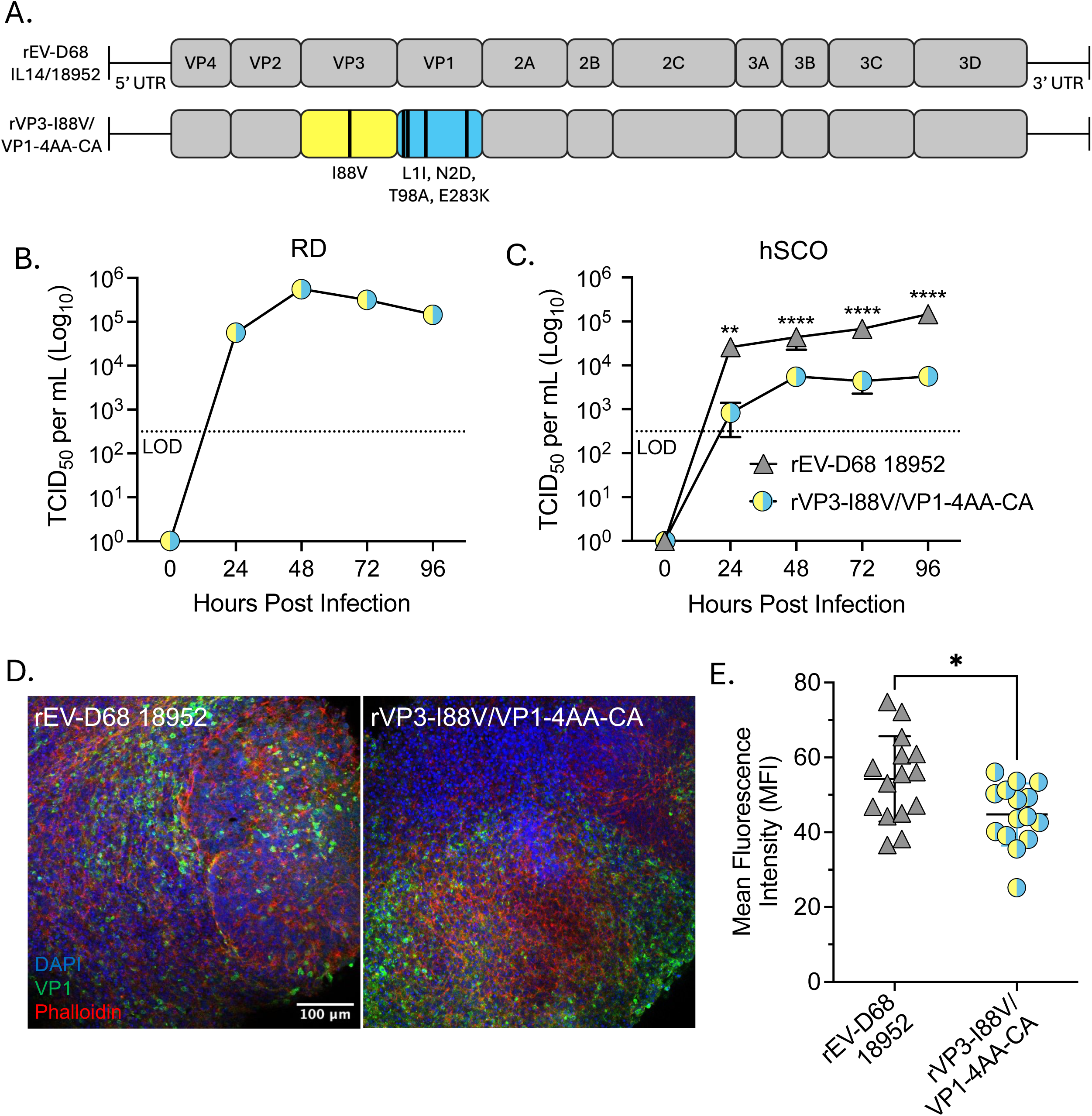
Combining VP1-4AA-CA and VP3-I88V-MO point mutations results in decreased viral titer compared to rEV-D68 18952. A.) Schema of rVP3-I88V/VP1-4AA-CA recombinant viruses with VP3 I88V and VP1 L1I/N2D/T98A/E283K. Gray represents rEV-D68 18952, and yellow and blue represent rVP3-I88V/VP1-4AA-CA. B.) RD cells were infected at an MOI 0.01 PFU/cell in biological triplicate with rVP3-I88V/VP1-4AA-CA. Viral titers were determined by TCID_50_ in RD cells. C.) SCTi003A hSCO were infected with 10^4^ PFU of either rEV-D68 18952 (gray triangle) or rVP3-I88V/VP1-4AA-CA (yellow and blue circle) in biological triplicate. Supernatants were collected every 24 hours up to 96 hpi. Viral titers were determined by TCID50 in RD cells. A two-way ANOVA with multiple comparisons to EV-D68 18952 was performed (* p < 0.05 ** < 0.005 **** < 0.00005). D. IF images of SCTi003A hSCOs infected with rEV-D68 18952 or rVP3-I88V/VP1-4AA-CA. hSCOs were infected with 10^4^ PFU and were fixed in 4% paraformaldehyde at 96 hpi. hSCOs were then stained with DAPI (blue), VP1 (green), and phalloidin (red). E.) MFI of VP1 staining was quantified across five biological replicate images of each recombinant virus across multiple Z stacks and organoids. Quantified values were calculated using Fiji ImageJ. An unpaired student t-test was performed (* p < 0.05).

We then infected hSCOs at 14 days post-differentiation with 10^4^ PFU of rVP3-I88V/VP1-4AA-CA and rEV-D68 18952 and assessed viral titer and VP1 expression and localization with IF at 96 hpi (Figure 3C-E). Compared to rEV-D68 18952, we found that rVP3-I88V/VP1-4AA-CA viral titer and VP1 expression was significantly reduced at each evaluated time point until 96 hpi.

### Substitution frequency of EV-D68 VP1 and VP3 residues of interest since 2014

In 2014, the CDC began tracking AFM cases coincident with EV-D68 circulation. Infections continued biannually until 2020 when there was minimal EV-D68 circulation due to non-pharmaceutical interventions for COVID-19 [10, 11]. While EV-D68 returned robustly in in 2022, there was not a corresponding spike in AFM cases [12, 13]. Therefore, we sought to analyze available sequencing data to assess if changes in the neurotropism driving residues of VP3 and VP1 have changed at the population level provided a potential explanation for the altered clinical neurotropism phenotype seen since 2022. We assessed 1,968 sequences from strains isolated between 2014-2024 worldwide. We found that since 2014, VP3 I88, the virulent residue, has been fixed at the population level with only three clinical isolates (2016-2018), with VP3 V88 (Figure 4A). As for VP1, we found that E283 is very stable with only seven occurrences of K283 since 2014. The other VP1 sites of interest were more variable, however, with I1 occurring in 61% of sequences, D2 occurring in 59%, and A98 occurring in 43%, potentially explaining the changing clinical manifestation. Some substitutions resulted in dramatic shifts in amino acid side chain properties as well, with VP1 T98 shifting from polar to non-polar (Figure 4C).

**Figure 4.**
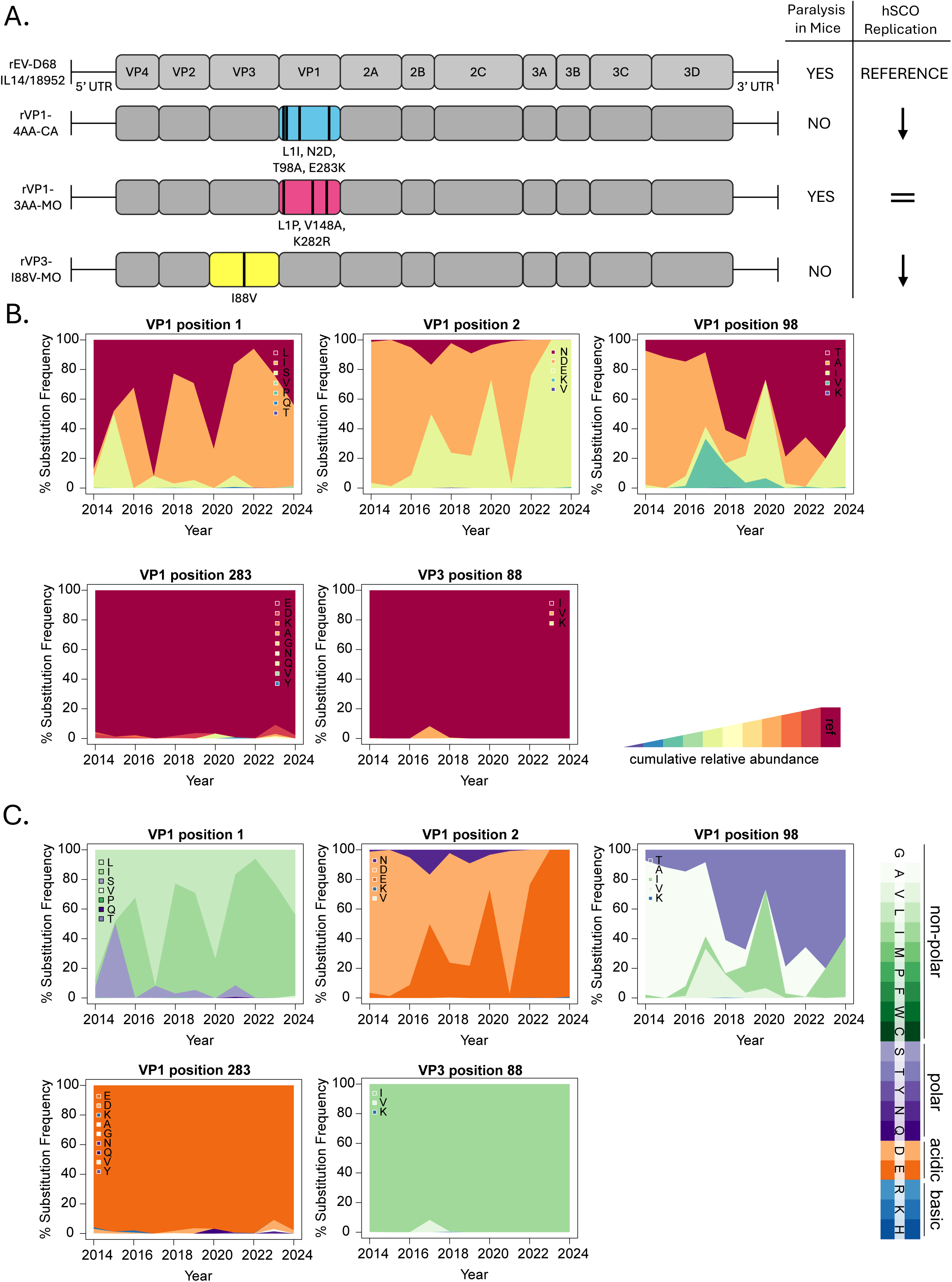
Frequency of VP1 and VP3 point mutations in clinical isolates that have circulated between 2014 to 2024. A. Schema of VP1 and VP3 recombinant viruses and the effect they have on mice paralysis and on hSCO replication. Gray represents rEV-D68 18952, blue – rVP1-4AA-CA (L1I/N2D/T98A/E283K), pink – rVP1-3AA-MO (L1P/V148A/K282R), and yellow – rVP3-I88V-MO. Black arrow – decreased viral replication in hSCO compared to reference strain (EV-D68 18952), black ‘=’ – similar rate of viral replication to EV-D68 18952. B. Global EV-D68 sequences from 2014-2024 were obtained from the Bacterial and Viral Bioinformatics Resource Center (n=1,968) and aligned using NextClade. Amino acid substitutions were visualized in an area plot as a percentage of the total sequences analyzed in each year. The identity of each residue in the reference strain, EV-D68 18952, is shown in red, and substitutions are colored according to relative abundance in each year. C. Amino acid types were overlaid onto the substitution frequencies shown in panel B.

## DISCUSSION

EV-D68 is a known cause of AFM thought to be driven by viral replication in the spinal cord. Previous research in immunocompromised murine models has demonstrated that EV-D68 containing point mutations in structural proteins VP3 and VP1 resulted in decreased paralysis and less morbidity [3, 4]. We sought to investigate if VP3 and VP1 directed neurotropism in a human CNS model, if those attenuating mutations were additive, and if these mutations persisted in modern EV-D68 clinical sequences, perhaps accounting for the decrease in AFM associated with EV-D68 since 2022.

In this report, we investigated EV-D68 neurotropism determinants with a multicellular human CNS model. We found that rVP3-I88V-MO and rVP1-4AA-CA resulted in decreased infection in hSCOs whereas rVP1-3AA-MO did not (Figure 2). These data recapitulate studies from both Yeh et al. and Leser et al., highlighting the use of hSCOs as a complementary approach to study enterovirus infection in the context of the human CNS. We then found that combining VP3 I88V and VP1 L1I/N2D/T98A/E283K into a single recombinant virus (rVP3-I88V/VP1-4AA-CA) resulted in decreased viral titer and VP1 staining in hSCOs compared to rEV-D68 18952 (Figure 3). However, when rVP3-I88V/VP1-4AA-CA was compared to rVP3-I88V-MO and rVP1-4AA-CA (Figure 2C), they seem to achieve similar viral titers over time (Figure 2C and 3C), suggesting that these mutations are not additive.

Each of these prior studies used different inoculation routes, with Yeh et al. using both IC and IP injection and Leser et al. using intramuscular (IM) injection [3, 4]. Viruses that are injected by IM and IP enter the CNS indirectly by moving into nearby blood vessels to enter the blood stream [14, 15]. Viruses injected by IM or IP then must cross the blood brain barrier (BBB) or exploit retrograde axonal transport to move from the peripheral nerves directly to motor neurons within the spinal cord [14, 15]. IC injection, however, delivers viral infection directly to the CNS, bypassing the BBB. Similarly, infection of hSCOs mirrors direct inoculation of the CNS, allowing for the study of tropism. Leser et al. did not assess IC infection or infection of neurons *in vitro*, so it is not known if the infection defect is due to an issue in cellular tropism or a defect of trafficking to and accessing the CNS. They also did not assess each mutation individually; thus, it is unknown which mutation or combination of mutations are required to modify EV-D68 neurovirulence. Our data in hSCO suggests that this virus, rVP1-4AA-CA, has replication at least partially limited by tropism.

In 2014, the CDC began tracking clinical sequences of EV-D68 and its association to AFM, however, in 2022, AFM associated EV-D68 cases decreased [12, 13]. The reasons for this decoupling are unknown. Yeh et al. determined the prevalence of VP3-I88V in clinical isolates from the former Virus Pathogen Resource (ViPR) from 2014 [3]. Consistently, their analysis found that only two out of 676 collected sequences had V88 present in VP3. Our analysis of an additional 1,292 sequences from the recently consolidated Bacterial and Viral Bioinformatics Resource Center (BV-BRC) found only one more virus with V88. Past and present circulating clinical isolates of EV-D68 have VP3 I88 as the prominent amino acid (Figure 4B). We therefore anticipate that it is unlikely that I88V is the main driver of decreased neurovirulence in recent clinical EV-D68 strains.

Another comprehensive examination of EV-D68 sequences in 2023 (published in 2025) investigated European clinical samples and compared them to Leser et al. findings [4, 16]. They found that A98T was found in all European sequences while E283K was identified in all except two. Finally, they identified that the VP1 N2D substitution has continued to evolve into D2E. In 2022, the Johns Hopkins Health System also identified a major shift to VP1 E2 which has remained the primary circulating residue in isolates from Baltimore, MD [17]. Similarly, we found that compared to reference strain EV-D68 18952, there are only seven occurrences of K283 since 2014, indicating that this mutation is prominent not only in European isolates, but globally. We anticipate that VP1 site 283 is not a primary driver of neuropathogenesis in recent circulating strains due to the limited variability in amino acids (Figure 4B and 4C). By contrast, VP1 sites 1, 2, and 98 have a greater percent substitution frequency between 2014 and 2024. I1 occurred in 61% of sequences, and A98 occurred in 43% (Figure 4B). Like Hirvoven et al., we found that D2 occurred in 59% of isolates and have later shifted to an E (39% relative abundance). These data indicate that there is selective pressure on VP1 at these specific sites. VP1 site 2 has evolved from a basic amino acid (N) to primarily acidic amino acids (D and E). Interestingly, D and E are biochemically similar, differing only by an extra methylene in the glutamic acid side chain. VP1 site 98 similarly has evolved from a polar amino acid (T) to different types of non-polar amino acids (A, I, and V). Perhaps changes in amino acid charge and solubility have altered enzymatic active sites, substrate and receptor binding, and protein-protein interactions, rendering recent strains to be non-neuropathogenic.

Additionally, both VP3 and VP1 are structural proteins that form the viral capsid with VP2 while VP4 is nestled within the particle to stabilize the structure. VP1 also functions by mediating viral receptor binding, host immune evasion, regulation of tissue tropism, and encodes epitopes that provide the basis for EV-D68 serotyping [18-20]. Structurally, VP1 also has two surface exposed loops, the BC-loop and DE-loop, both of which interact with host cell receptors and can modulate receptor binding affinity [19]. For example, host receptor major facilitator superfamily domain-containing 6 (MFSD6) interactions are driven by the flexibility of the BC-loop [21]. Because the BC-loop is comprised of amino acids between positions 90-103 of VP1 [22], there is a possibility that VP1 site 98 helps drive host cell receptor interactions and may account for decreased AFM in the 2022 and 2024 seasons.

In this study, we evaluated previously described neurovirulence determinants in structural proteins VP3 and VP1 to investigate neurotropism of EV-D68 in hSCO. Comprehensive and systematic evaluation of neurovirulence drivers of other structural and non-structural proteins will provide additional insight into the clinical manifestations of this continually evolving pathogen.

### Conclusions

In this study, we evaluated previously described EV-D68 neurovirulence determinants in a multicellular human CNS model. Infection of hSCOs with recombinant viruses resulted in similar replication results as previously published murine models, further demonstrating model complementarity. We found that combining VP1 L1I/N2D/T98A/E283K and VP3 I88V mutations did not result in an additive or abortive phenotype in hSCOs. We then evaluated the frequency of these point mutations in reported clinical EV-D68 isolates since 2014 and discovered significant ongoing evolution at three sites: VP1 1, 2, and 98, perhaps explaining changes in clinical phenotype and lessened neurovirulence. These data show that EV-D68 continues to rapidly adapt and evolve, thus emphasizing the importance for continued EV-D68 surveillance and study of contemporary strains.

## Author Contributions

Conceptualization, M.C.F and J.E.P.; Methodology, M.C.F. and J.E.P; Validation, J.E.P.; Data Curation, J.E.P., G.Y., and J.E.J.; Writing – Original Draft Preparation, J.E.P.; Writing – Review & Editing, M.C.F., J.E.P., and J.E.J; Visualization, J.E.P. and J.E.J.; Supervision, M.C.F.; Funding Acquisition, M.C.F.

## Funding

M.C.F., J.E.P., J.E.J, and G.Y. were supported by NIH NIAID K08AI171177, NIH NIAID R21AI193384, and the Richard King Mellon Foundation.

## Institutional Review Board Statement

Not applicable.

## Informed Review Board Statement

Not applicable.

## Data Availability Statement

Not applicable.

## Acknowledgements

We thank the members of the Freeman and Eddens lab for many useful discussions about this work. Model figures were prepared using BioRender.com. Confocal images were captured in the Cell Imaging Core Facility at the UPMC Children’s Hospital of Pittsburgh Rangos Research Center. We are grateful to the Cell Imaging Core staff for their technical assistance.

## Conflicts of Interest

The authors declare no conflicts of interest.

## Notes

### Competing Interest Statement

The authors have declared no competing interest.

## Bibliography

1. Schieble, J.H., V.L. Fox, and E.H. Lennette, A probable new human picornavirus associated with respiratory diseases. Am J Epidemiol, 1967. 85(2): p. 297–310.

2. Messacar, K. and K.L. Tyler, Enterovirus D68-Associated Acute Flaccid Myelitis: Rising to the Clinical and Research Challenges. Jama, 2019. 321(9): p. 831–832.

3. Yeh, M.T., et al., Mapping Attenuation Determinants in Enterovirus-D68. Viruses, 2020. 12(8).

4. Leser, J.S., et al., VP1 is the primary determinant of neuropathogenesis in a mouse model of enterovirus D68 acute flaccid myelitis. J Virol, 2024. 98(7): p. e0039724.

5. Aguglia, G., et al., Contemporary enterovirus-D68 isolates infect human spinal cord organoids. mBio, 2023. 14(4): p. e0105823.

6. Jones, J.E., et al., Recombinant B3 clade enterovirus D68 strains are efficiently rescued in 293T cells and infect human spinal cord organoids. bioRxiv, 2024: p. 2024.12.20.629498.

7. Oberste, M.S., et al., Improved molecular identification of enteroviruses by RT-PCR and amplicon sequencing. J Clin Virol, 2003. 26(3): p. 375–7.

8. Ivan Aksamentov, C.R., Emma B Hodcroft, Richard A. Neher, Nextclade: clade assignment, mutation calling and quality control for viral genomes. 2021, Zenodo: Journal of Open Source Software.

9. Neuner-Jehle, N., González-Sánchez, Alejandra, Hodcroft, Emma, and European Non-Polio Enterovirus Network (ENPEN), enterovirus-phylo/nextclade_d68: Enterovirus D68 Nextclade Dataset v1.0.1. 2025, Zenodo.

10. Melisa M. Shah, M.A.P., MPH; Joana Y. Lively, MPH; Vasanthi Avadhanula, PhD; Julie A. Boom, MD; James Chappell, MD, PhD; Janet A. Englund, MD; Wende Fregoe; Natasha B. Halasa, MD; Christopher J. Harrison, MD; Robert W. Hickey, MD; Eileen J. Klein, MD; Monica M. McNeal, MS; Marian G. Michaels, MD; Mary E. Moffatt, MD; Catherine Otten, MD; Leila C. Sahni, PhD; Elizabeth Schlaudecker, MD; Jennifer E. Schuster, MD; Rangaraj Selvarangan, PhD; Mary A. Staat, MD; Laura S. Stewart, PhD; Geoffrey A. Weinberg, MD; John V. Williams, MD; Terry Fan Fei Ng; Janell A. Routh, MD; Susan I. Gerber, MD; Meredith L. McMorrow, MD; Brian Rha, MD; Claire M. Midgley, PhD, Enterovirus D68-Associated Acute Respiratory Illness ─ New Vaccine Surveillance Network, United States, July–November 2018–2020. 2021: Morbidity and Mortality Weekly Report (MMWR).

11. Sonja J. Olsen, P.A.K.W., MPH; Alicia P. Budd, MPH; Mila M. Prill, MSPH; John Steel, PhD; Claire M. Midgley, PhD; Krista Kniss, MPH; Erin Burns; Thomas Rowe, MS; Angela Foust; Gabriela Jasso; Angiezel Merced-Morales, MPH; C. Todd Davis, PhD; Yunho Jang, PhD; Joyce Jones, MS; Peter Daly, MPH; Larisa Gubareva, PhD; John Barnes, PhD; Rebecca Kondor, PhD; Wendy Sessions, MPH; Catherine Smith, MS; David E. Wentworth, PhD; Shikha Garg, MD; Fiona P. Havers, MD; Alicia M. Fry, MD; Aron J. Hall, DVM; Lynnette Brammer, MPH; Benjamin J. Silk, PhD, Changes in Influenza and Other Resporatory Virus Activity During the COVID-19 Pandemic - United States, 2020-2021. 2021: Morbidity and Mortality Weekly Report (MMWR).

12. AFM Cases and Outbreaks. 2025 December 3, 2025 [cited 2025 December 11]; Available from: https://www.cdc.gov/acute-flaccid-myelitis/cases/index.html.

13. Kevin C. Ma, P.A.W., MPH; Heidi L. Moline, MD; Heather M. Scobie, PhD; Claire M. Midgley, PhD; Hannah L. Kirking, MD; Jennifer Adjemian, PhD; Kathleen P. Hartnett, PhD; Dylan Johns, MS; Jefferson M. Jones, MD; Adriana Lopez, MHS; Xiaoyan Lu, MS; Ariana Perez, MPH; Cria G. Perrine, PhD; Andzelika E. Rzucidlo, MPH; Meredith L. McMorrow, MD; Benjamin J. Silk, PhD; Zachary Stein, MPH; Everardo Vega, PhD; New Vaccine Surveillance Network Collaborators; Aron J. Hall, DVM1, Increase in Acute Respiratory Illnesses Among Children and Adolescents Associated with Rhinoviruses and Enteroviruses, Including Enterovirus D68 — United States, July–September 2022. Morbidity and Mortality Weekly Report, 2022.

14. Tosolini, A.P. and J.N. Sleigh, Intramuscular Delivery of Gene Therapy for Targeting the Nervous System. Frontiers in Molecular Neuroscience, 2020. Volume 13 - 2020.

15. Salameh, T.S. and W.A. Banks, Delivery of therapeutic peptides and proteins to the CNS. Adv Pharmacol, 2014. 71: p. 277–99.

16. Hirvonen, A., et al., Sustained circulation of enterovirus D68 in Europe in 2023 and the continued evolution of enterovirus D68 B3-lineages associated with distinct amino acid substitutions in VP1 protein. Journal of Clinical Virology, 2025. 178: p. 105785.

17. Fall, A., et al., An increase in enterovirus D68 circulation and viral evolution during a period of increased influenza like illness, The Johns Hopkins Health System, USA, 2022. Journal of Clinical Virology, 2023. 160: p. 105379.

18. Lin, X., et al., A Novel Peptide from VP1 of EV-D68 Exhibits Broad-Spectrum Antiviral Activity Against Human Enteroviruses. Biomolecules, 2024. 14(10): p. 1331.

19. Liu, Y., et al., Structure and inhibition of EV-D68, a virus that causes respiratory illness in children. Science, 2015. 347(6217): p. 71-4.

20. Zhu, Y., L. Wang, and J. Shen, Enterovirus D68 Sequence Variations and Pathogenicity: A Review. Viruses, 2026. 18(1): p. 73.

21. Liu, X., et al., MFSD6 is an entry receptor for respiratory enterovirus D68. Cell Host & Microbe, 2025. 33(2): p. 267–278.e4.

22. Shi, Y., et al., Molecular epidemiology and recombination of enterovirus D68 in China. Infection, Genetics and Evolution, 2023. 115: p. 105512.

